# Network-based prediction approach for cancer-specific driver missense mutations using a graph neural network

**DOI:** 10.1101/2023.07.05.547896

**Authors:** Narumi Hatano, Mayumi Kamada, Ryosuke Kojima, Yasushi Okuno

**Affiliations:** Graduate School of Medicine and Faculty of Medicine, Kyoto University, Kyoto, Japan; RIKEN Center for Computational Science(R-CCS)HPC/HPC- and AI-driven Drug Development Platform Division, Kobe, Japan

**Keywords:** Driver mutation prediction, Cancer missense mutation, Graph Neural Network, Molecular interaction

## Abstract

**Background:** In cancer genomic medicine, finding driver mutations involved in cancer development and tumor growth is crucial. Machine-learning methods to predict driver missense mutations have been developed because variants are frequently detected by genomic sequencing. However, even though the abnormalities in molecular networks are associated with cancer, many of these methods focus on individual variants and do not consider molecular networks. Here we propose a new network-based method, Net-DMPred, to predict driver missense mutations considering molecular networks. Net-DMPred consists of the graph part and the prediction part. In the graph part, molecular networks are learned by a graph neural network (GNN). The prediction part learns whether variants are driver variants using features of individual variants combined with the graph features learned in the graph part.

**Results:** Net-DMPred, which considers molecular networks, performed better than conventional methods. Furthermore, the prediction performance differed by the molecular network structure used in learning, suggesting that it is important to consider not only the local network related to cancer but also the large-scale network in living organisms.

**Conclusions:** We propose a network-based machine learning method, Net-DMPred, for predicting cancer driver missense mutations. Our method enables us to consider the entire graph architecture representing the molecular network because it uses GNN. Net-DMPred is expected to detect driver mutations from a lot of missense mutations that are not known to be associated with cancer.

## Background

Genomic sequencing studies have been a massive advancement in cancer genomic medicine. In conventional cancer medicine, treatment is uniformly determined by characteristics such as anatomical site and progression stage. However, some patients do not respond well to treatment. Cancer genomic medicine has become popular because gene mutations are associated with cancer development. In this medicine, the best treatment is selected based on the genomic background of each patient. For this reason, cancer genomic medicine is expected to enhance therapeutic effects and reduce side effects. One of the challenges in cancer genomic medicine is the clinical interpretation of variants. Although a lot of variants are detected by genomic analysis, most of them are passenger mutations not directly involved in cancer development. A small fraction of variants are driver mutations that are involved in cancer development [1]. Therefore, it is important to distinguish between driver mutations and passenger mutations.

It is time-consuming and expensive to validate whether variants are driver mutations, so machine-learning methods have been developed to predict if missense mutations are driver mutations. For example, CHASM [2] and CHASMplus [3] predict driver mutations by utilizing features obtained from conserved sequences and protein structure to characterize each variant. CanDrA [4] is an ensemble tool that uses the results of other prediction tools as variant features.

In living organisms, molecular networks are formed by various molecular interactions and signal transduction pathways, and abnormalities in molecular networks are associated with cancer. However, despite their importance, most existing tools for driver mutation prediction do not consider molecular networks. Although CHASMplus uses the number of molecular interactions for the amino acid site of a mutation as a variant feature, it only considers local molecular relationships and does not consider molecular networks. Molecular networks can be represented as a graph with molecules as nodes and molecular relationships as edges. Some prediction methods utilizing a graph to consider molecular interactions have been developed. Network&AA [5] is a driver prediction tool that considers the centrality of graphs representing molecular networks. However, the previous methods only consider one aggregated aspect of the graph, such as centrality, and not the entire graph structure.

Here we proposed a new network-based prediction method, Net-DMPred. This method uses a graph neural network (GNN) and learns a graph represented as molecular networks. GNN is a deep learning method for graphs and can learn the entire graphical structure. Net-DMPred predicts driver mutations using the features of the molecular networks by GNN as background knowledge and combining them with the features of individual variants used in conventional methods.

## Results and Discussion

### Overview of Net-DMPred

Net-DMPred consists of the graph and prediction part (Figure 1). The graph part learns background knowledge and the prediction part learns combining individual information and background knowledge learned by the graph part. In this study, molecular networks were used as background knowledge and individual variant features as individual information.

**Fig. 1.**
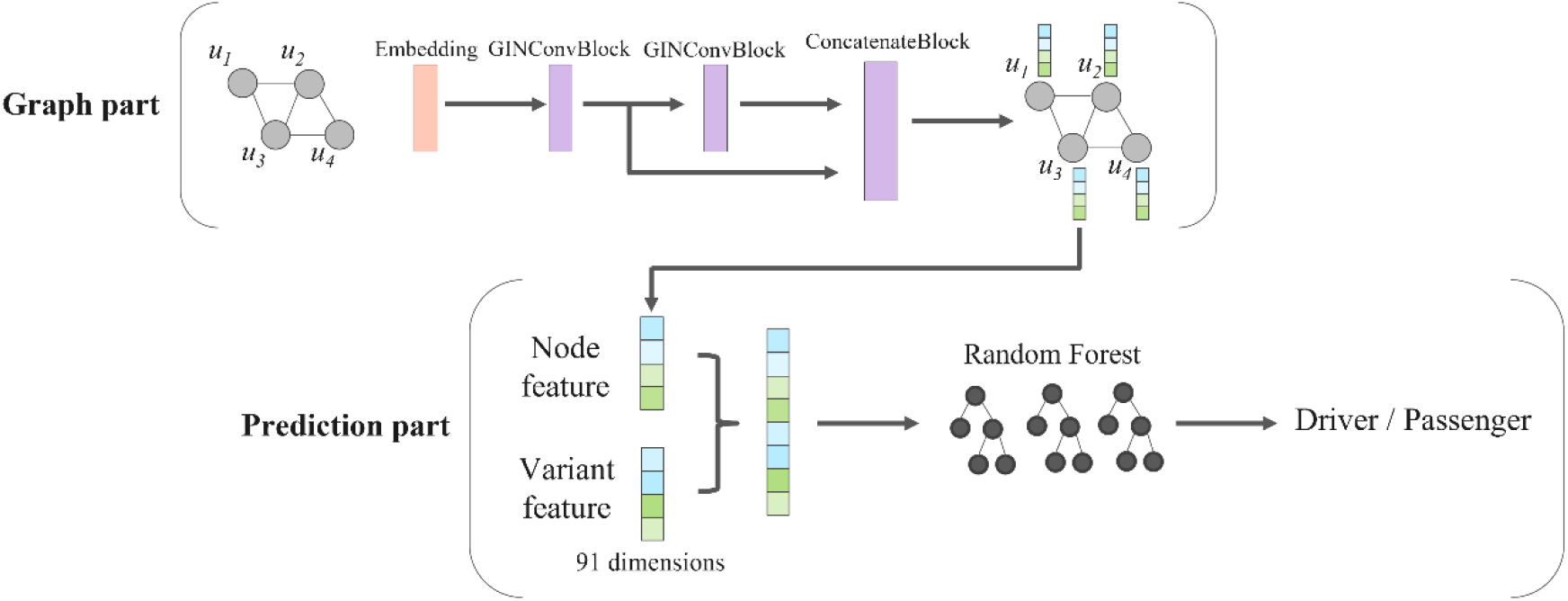
Model Architecture.

In the graph part, graphs representing molecular networks are learned using GNN, and feature vectors for each molecular node are computed. GNN can consider the entire architecture of graph. In the prediction part, the variants are predicted to be driver or passenger by Random Forest with individual variant features and graph node features corresponding to the genes with variants. This framework enables us to utilize general molecular networks learned in the graph part as common background knowledge of individual variants in predicting driver mutation.

### Performance evaluation of the training dataset

We used the training dataset of gene mutations provided by CHASMplus (http://karchinlab.org/data/CHASMplus/formatted_training_list.txt.gz, acquired on October 28, 2021). We performed the under sampling on this training dataset, and 928 driver and 3,712 passenger mutations were used for training.

As features of gene mutations, we used 91 features obtained from SNVBox database [6] and the outputs of HotMAPS 1D method [7] following CHASMplus.

To investigate the relationship between the graph architecture and the prediction performance, we constructed three knowledge graphs representing molecular networks, “Cancer pathway,” “Molecular interaction,” and “Cancer pathway + Molecular interaction” (Figure 2, Table 1). “Cancer pathway” was obtained from PathwayMapper [8] (acquired on November 1, 2021). PathwayMapper contains 10 public pathways [9] (such as TP53 pathway and the PI3K pathway) known to be associated with pan-cancer, and this study used these 10 pathways. “Molecular interaction” was obtained from Pathway Commons v12 [10]. Pathway Commons is public pathway and molecular interaction databases. “Cancer pathway + Molecular interaction” was combining “Cancer pathway” and “Molecular interaction”. Moreover, to confirm the usefulness of our framework, we performed the prediction using only individual variant features without the graph features (hereinafter referred to as “No graph”). It corresponds to the conventional prediction approach.

**Fig. 2.**
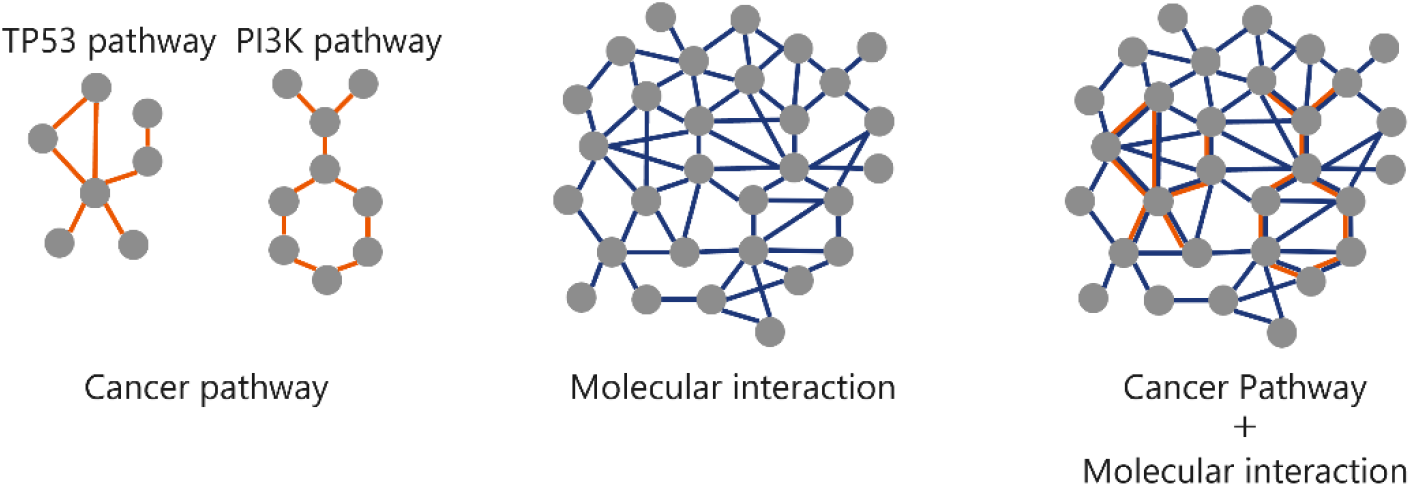
Three knowledge graphs. “Cancer pathway” includes 10 pathways such as the TP53 pathway and the PI3K pathway.

**Table 1.** Details of three knowledge graphs

| Graph | Node | Edge | Edge type |
| --- | --- | --- | --- |
| Cancer pathway | 200 | 870 | 23 |
| Molecular interaction | 30899 | 3672040 | 13 |
| Cancer pathway + Molecular interaction | 30923 | 3672910 | 36 |

Number of nodes, number of edges, and types of edges in the three graphs.

The models were evaluated the mean of ROC-AUC (Area Under the Receiver Operatorating Characteristic Curve) and PR-AUC (Area Under the Precision-Recall Curve) with five-fold cross-validation. In this study, the dimension of the graph node vector was 16 and 32.

Figure 3 shows the results of the performance evaluation of the training dataset. Comparing the mean value of ROC-AUC and PR-AUC, the models with graphs, “Cancer pathway,” “Molecular interaction,” and “Cancer pathway + Molecular interaction” performed better than the model without a graph, “No graph.” These results show that considering molecular networks is useful for driver mutation prediction.

**Fig. 3.**
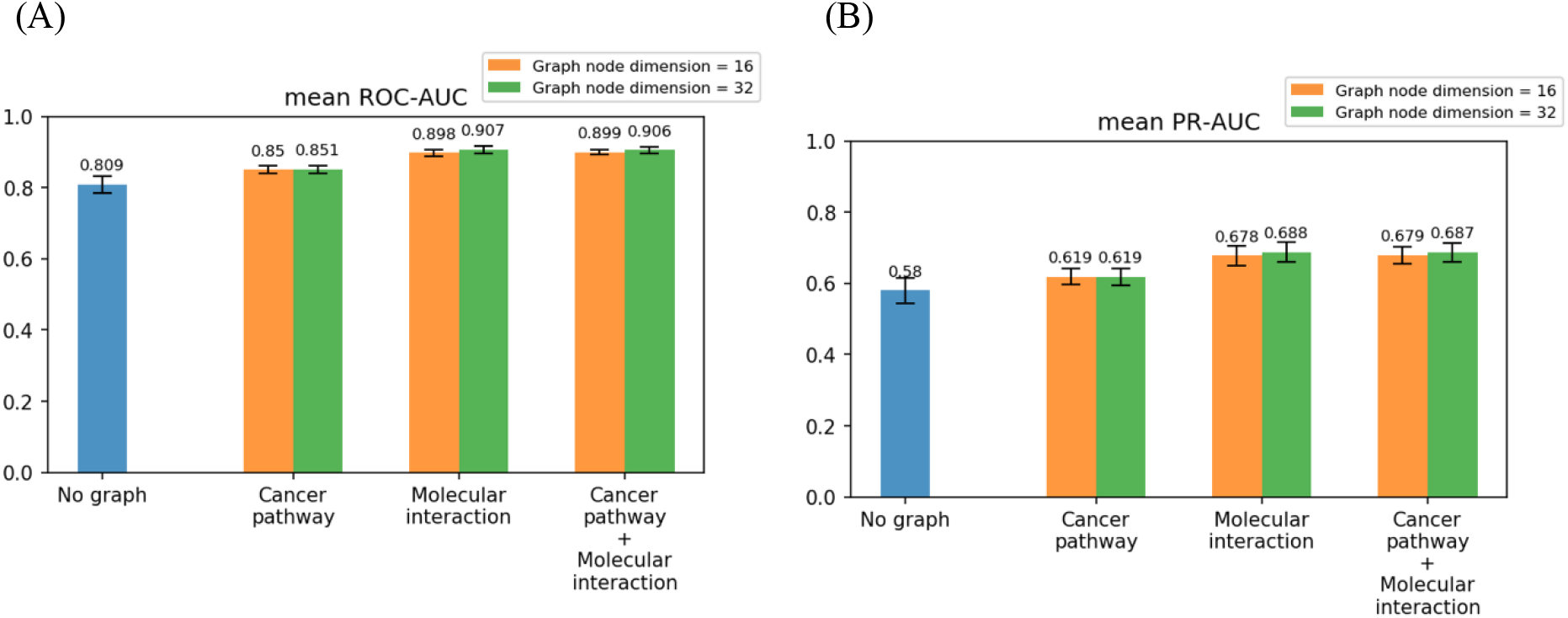
Results of performance evaluation of the training dataset. (A) Mean value of ROC-AUC (B) Mean value of PR-AUC

Comparing the differences in graph architectures, “Molecular interaction” and “Cancer pathway + Molecular interaction” performed better than “Cancer pathway.” This result demonstrates that the prediction performances were increased when not only the local network, “Cancer pathway,” but also the large-scale network, “Molecular interaction,” were used as molecular networks. It also indicates that it is important to construct appropriate graphs because the prediction performances differed by graphs.

For the dimension of the node vectors, the prediction performance was slightly higher when 32, rather than 16 dimensions, were used. Therefore, we used the node features with 32 dimensions in the following analysis.

### Performance evaluation with the benchmark datasets

We compared our approach with existing methods using five benchmark datasets obtained from Tokheim and Karchin [3] http://karchinlab.org/data/CHASMplus/Tokheim_Cell_Systems_2019.tar.gz, acquired on October 28, 2021); Kim *et al.* [11], IARC TP53 [12], Ng *et al.* [13], Gene panel (OncoKB) [14, 15], CGC-recurrent [16]. These datasets were derived from *in vivo* and *in vitro* experiments and literature. Each dataset had different criteria for positive (driver) and negative (passenger) mutations. Therefore, these five datasets allowed for the multifaceted evaluation of the prediction model. In these datasets, some gene mutations overlapped with the training dataset. To strictly evaluate the prediction model, gene mutations that overlapped with the training dataset were dropped from benchmark datasets. Table 2 shows counts of gene mutation and unique genes with mutations used in benchmark datasets.

**Table 2.** Benchmark datasets

| Benchmark | Data source | Positive | Negative | Protein unique |
| --- | --- | --- | --- | --- |
| Kim et al. | *in vivo* | 8 | 17 | 11 |
| IARC TP53 | *in vitro* | 375 | 1533 | 1 |
| Ng et al. | *in vitro* | 116 | 357 | 43 |
| Gene panel (OncoKB) | Literature-based | 329 | 38596 | 410 |
| CGC-recurrent | Literature-based | 33 | 6805 | 3261 |

Number of positive and negative data and unique of proteins with mutations.

Figure 4 shows the prediction performances of the proposed models and the existing prediction tools for cancer driver mutations; CHASM, TransFIC [17], CanDrA, ParsSNP [18], CHASMplus, Network&AA. (The performances of other tools are shown in Additional file 1-2: Supplementary Table 1–2.) The prediction performance of our models with the graph, “Cancer pathway,” “Molecular interaction,” and “Cancer pathway + Molecular interaction,” was better or comparable to the model without a graph, “No graph.”

**Fig. 4.**
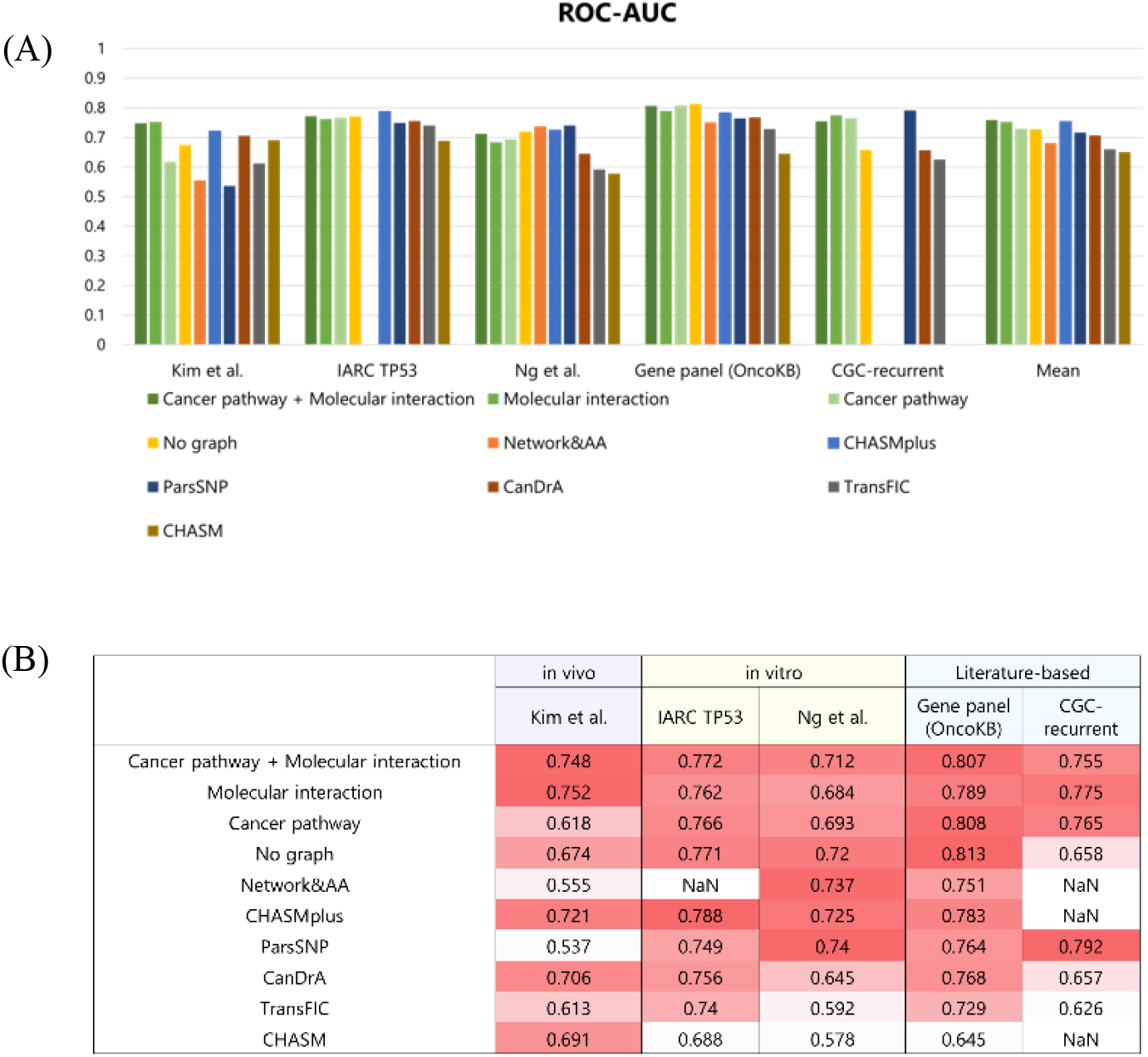
Results of five benchmark datasets. (A) Bar graph of ROC-AUC score. (B) Table of ROC-AUC score. In Network&AA, we could not acquire prediction scores of variants in IARC TP53 and CGC-recurrent. In CHASMplus and CHASM, variants in CGC-recurrent did not have prediction scores.

The proposed models showed higher performance than Network&AA, which considers the centralities of molecular networks. These results show that it is important to consider the entire molecular networks.

Moreover, in the proposed models, “Molecular interaction” and “Cancer pathway + Molecular interaction” showed overall higher performances than “Cancer pathway.” These results show that the prediction performances were increased when not only the local network, “Cancer pathway,” but also the large-scale network, “Molecular interaction,” were used. The performance on some benchmark datasets was increased when the models with graphs, but others were not increased. It may be caused by the variety and differences between the five benchmark datasets in terms of the label definition and the data source.

For the Kim et al. dataset, the models with graphs showed higher performance than conventional tools. Kim et al. dataset was derived from in vivo and evaluated for the impact of tumor growth on missense mutation in mice. It can be speculated that the consideration of molecular networks may be useful in predicting in vivo datasets, such as this data.

On the other hand, in the prediction of IARC TP53 and Ng et al. datasets, the prediction performances of the models with graphs were not increased. IARC TP53 and Ng et al. datasets were derived from in vitro. The criteria for driver and passenger were transactivation levels for TP53 targets in IARC TP53 dataset and cell viability in Ng et al. dataset, respectively; these criteria differed from those in the training dataset.

The prediction performance of our models for the Gene panel (OncoKB) dataset was not improved. This dataset uses OncoKB annotations as labels. Mutations annotated with “Oncogenic” and “Likely Oncogenic” were defined as positive, and mutations with other annotations were defined as negative. In the negative data, there were missense mutations not only annotated with “Likely Neutral” and “Inconclusive” but also “Unknown” in OncoKB. Therefore, there is the possibility that some missense mutations which were labeled as negative in the dataset may be positive. Among 38,925 mutations in the Gene panel (OncoKB) dataset, the proposed model “Cancer pathway + Molecular interaction” predicted 8,876 mutations as the driver mutations, 8,624 of which were annotated as “Unknown” in the provided Gene panel (OncoKB) dataset. Then, we confirmed the latest annotations for these 8,624 mutations. As of March 20, 2023, in the OncoKB database, 12 mutations (RUNX1 p.D198N, PIK3CA p.D549N, KRAS p.D33E, PDGFRB p.N666K, ATM p.R3008C, MET p.H1094Y, PIK3CB p.A1048V, ERBB2 p.Q709L, BRAF p.S467L, BRAF p.N581I, MAP2K1 p.L177V, JAK3 p.R657W) were annotated as “Oncogenic” and 593 mutations as “Likely Oncogenic.” This finding implies that our proposed method is expected to identify driver mutations from a lot of mutations of uncertain significance. In addition, of the 8,876 mutations in the benchmark dataset that our model predicted as the driver mutations, 7,966 mutations remained classified as “Unknown” according to OncoKB as of March 20, 2023. Some of these mutations have potential to be confirmed as driver mutations through further experimental validation.

In the prediction of CGC-recurrent, the models with graphs performed better than models without graphs. This dataset includes a variety of genes with mutations. In a dataset consisting of such a large number and variety of genes, the graph features of each molecule learned in the graph part may be useful to increase the prediction performance.

### Interpretation of results

We used SHAP (Shapley Additive exPlanations) [19] to evaluate the contribution of graph features. Figure 5 shows the driver prediction scores obtained from “Cancer pathway + Molecular interaction” and “No graph,” and the contribution rate of graph features in “Cancer pathway + Molecular interaction” for the prediction of Kim et al. dataset. Here the graph feature importance rate was defined as the sum of SHAP values of graph features (32 features) divided by the sum of SHAP values of all features (123 features). The straight lines in Figure 5 show the thresholds for the driver and passenger of each model. The upper left plots show missense mutations which “Cancer pathway + Molecular interaction” predicted to be positive and “No graph” predicted to be negative. Three missense mutations (AKT1 p.E267G, KRAS p.D33E, AKT1 p.R370C) indicated by the arrows have positive labels as the correct answers. These three mutations were correctly predicted by “Cancer pathway + Molecular interaction” and incorrectly predicted by “No graph.” The graph feature importance rates for these variants show that the graph features had a significant impact on the predictions of these three mutations. In other words, by using the graph features, the model with the graph could have been predicted as positive for mutations correctly, which have been predicted as negative in models without the graph.

**Fig. 5.**
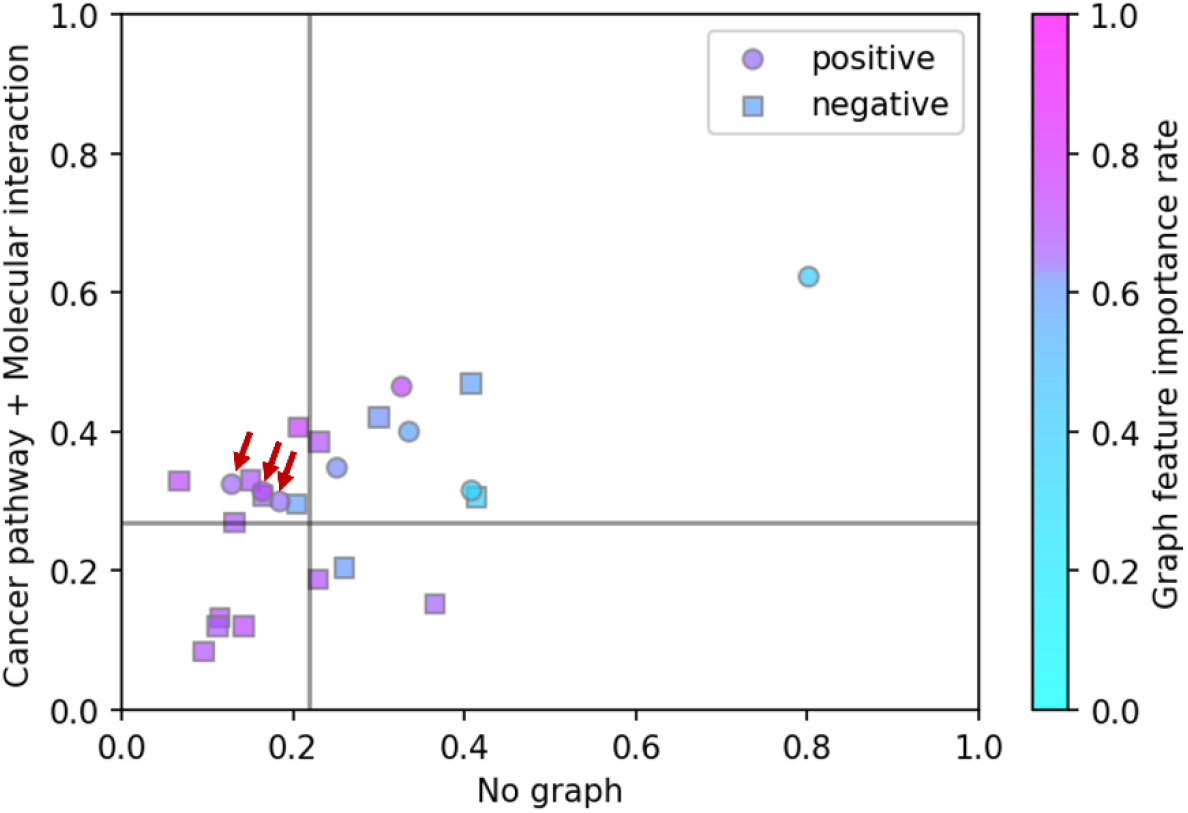
Interpretation of the results for Kim et al. dataset. This figure shows the prediction scores for “Cancer pathway + Molecular interaction” and “No graph,” and the graph feature importance rate for “Cancer pathway + Molecular interaction.” The circle plots are positive variants and square plots are negative variants in this benchmark dataset. The straight lines are thresholds for positive and negative for “Cancer pathway + Molecular interaction” and “No graph” (“Cancer pathway + Molecular interaction” = 0.270, “No graph” = 0.218). The plots with red arrows were predicted negative incorrectly for “No graph” but positive correctly for “Cancer pathway + Molecular interaction.”

## Conclusion

In this study, we proposed Net-DMPred, which predicts driver mutation considering the entire architecture of molecular networks. The performance of the proposed method showed higher than or comparable to the conventional methods. This result shows that it is important to have information about molecular networks in predicting cancer driver mutation.

The prediction performances differed by the graph architecture, considering not only the local network, “Cancer pathway,” but also the large-scale network, “Molecular interaction” improved the performance. This result indicates the importance of the construction of the graph appropriately.

We investigated the contribution of the graph to the prediction results using SHAP. It was confirmed that the graph features representing molecular networks contributed to the prediction of cancer driver mutation. However, the proposed method has limitations. The graph part, which learns the graph representing the molecular network, and the prediction part, which learns whether the mutation is driver or passenger, are independent of each other. Thus, we cannot interpret the contribution of each node in the graph, that is, related molecules in driver prediction.

Although the development of genome sequencing technology has facilitated the detection of variants, a lot of variants of uncertain significance have been accumulated. In this study, we demonstrated that Net-DMPred was able to predict the driver mutations that were previously unannotated but recently have been shown to be involved in cancer as the data accumulated. This result suggests that Net-DMPred holds promise in identifying driver mutations from a large number of mutations whose association with cancer is not yet known.

Our prediction model has room for improvement in two aspects. In the graph part, the performances were affected by graph architecture, and our proposed model can improve depending on the design of the graph. In the prediction part, while we used features such as amino acid properties, sequence conservation, and protein structure, our proposed model can utilize these features and various additional features such as cancer type and the result of conventional prediction tools.

## Methods

### Training dataset of gene mutation

The training dataset provided by CHASMplus consisted of 2,051 driver mutations and 616,515 passenger mutations from the missense mutation dataset based on The Cancer Genome Atlas (TCGA) [20]. Positive data (driver mutations) were defined as mutations that occurred in genes listed in Cancer Genome Landscapes [21] and with a lower mutation frequency (less than 500 mutations) in samples. Negative data (passenger mutations) were defined as other mutations.

This dataset was imbalanced in two respects; driver mutations were in a limited number of genes, and there were many more passenger mutations than driver mutations. We performed the following sampling steps on the training dataset to resolve these imbalances. First, driver and passenger mutations were sampled randomly for each gene up to the median of the driver mutations per gene (=21) to prevent mutations from being in limited genes. As a result, 928 driver mutations and 310,450 passenger mutations were obtained. Next, because 928 passenger mutations out of 310,450 occurred in genes with driver mutations, they were selected and always included in the training dataset. This process prevented excessive dependence on gene features on the dataset in predicting driver and passenger mutations. Finally, passenger mutations of genes that did not have driver mutations were randomly sampled in order that four times as many as driver mutations. As a result, the training dataset contained 4,640 mutations (928 driver mutations and 3,712 passenger mutations).

### Features of gene mutations

As the features of individual gene mutations, the proposed method uses 91 features. We obtained 88 features (Additional file 3: Supplementary Method) from SNVBox. It is a database of precomputed features for codons in the human exome, such as amino acid properties, sequence conservation, and protein structure. Other three features were obtained by running HotMAPS 1D. It is a method for estimating each gene’s recurrently mutated genome region (hotspot regions). In this study, we used gene mutation data extracted from TCGA as input data. We ran this method with window sizes (hyperparameters) of 0, 5, and 10. Then, we used the results (p-values) of the estimated regions as features of individual variants.

### Molecular network dataset

A knowledge graph representing molecular networks was constructed from two databases. The first was from molecular interaction data from Pathway Commons v12. The second was from cancer signaling pathway data from PathwayMapper. These datasets describe molecules and their relationships in living organisms.

Pathway Commons integrates more than 20 public pathways and interactions databases and describes relationships between proteins and small chemical compounds with 13 types of binary relationships. PathwayMapper is a web-based visualizing tool that includes various cancer-related pathways. Each cancer pathway contains information on regulatory relationships between molecules, such as activation and inhibition.

### Construction of knowledge graph

The molecular network dataset can be represented as a knowledge graph with molecules as nodes and molecular relationships as edges, and the type of molecular relationship can be represented as an edge label.

In this study, we constructed three knowledge graphs using the molecular networks and compared each graph’s difference in prediction performance. The first is the cancer pathway graph, “Cancer pathway,” obtained from PathwayMapper. The second is the molecular interaction graph, “Molecular interaction,” obtained from Pathway Commons. The third is the graph, “Cancer pathway + Molecular interaction,” combining “Cancer pathway” and “Molecular interaction.”

### Net-DMPred model architecture

#### Graph part

A graph consists of a pair of nodes and edges *G = (V, E)*. An edge is represented *(u, r, v)* ∈ E using node *u, v,* and the relation *r* where *u, v ∈ V, r∈ R*. *R* is a finite set of edge labels, *E* is a finite set of edges, and *V* is a finite set of nodes. In this study, background knowledge was molecular network, *V* was a set of molecules, and G was a set of molecular relationships.

In the graph part, the node vectors *z_u_* ∈ *R^C^* is calculated by the operation of the embedding layer and the graph neural network (GINConv) [22]:<colcnt=2>

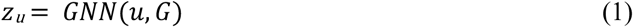

The architecture of this graph neural network has a connection between the output of each layer and the concatenate block to enhance expression of graph neural network. This graph neural network is constructed using a GIN block. A GIN block is defined as follows:

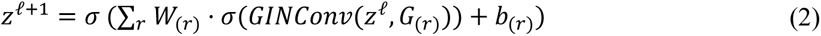

where z^ℓ^ represents a node vector of *ℓ*-th layer, *r∈R* is relation (edge label) in the graph *G*, and *G_(r)_* is defined as a subgraph that extracts the edges with relation *r* from the graph *G*.

The learning of the graph is pre-trained by link prediction. Link prediction is predicting the probability of the existence of an edge (link) between nodes. This pre-training learns node features based on the observed graph structure. The loss function is defined as follows:

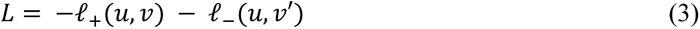

where *u, v* is randomly sampled from *E* and *v′* is randomly sampled from *V*.

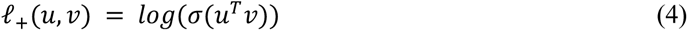

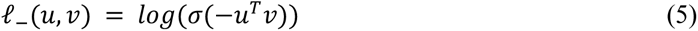

When a link with a relation is predicted, weight matrix is used:

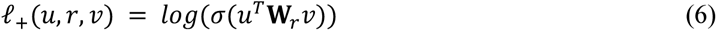

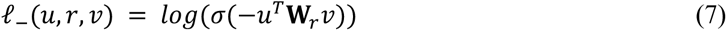

where ***W*** is a diagonal matrix.

In this study, the dimension of the node vector was *C* = 16 and 32, and the continuous random values were initially assigned to each vector. We employed a learning rate of 0.0001, and the model after 50 epochs was used for the analysis.

#### Prediction part

In the prediction part, the driver mutation prediction is performed using the features of individual variants combined with graph node features learned by the graph part. We used the Random Forest model [23] for prediction.

The probability of driver mutation *ŷ* is calculated using graph features and variant features as follows:

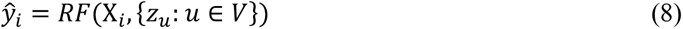

where *X_i_* is an variant feature of mutation *i* of gene *u*, and *z_u_* is a graph feature of gene *u*.

We performed Random Forest using the scikit-learn package (version 0.24.2). The hyperparameters of the model were tuned by three-fold grid search. Missing values were complemented by the mean.

### Performance evaluation with the benchmark datasets

To compare our approach with existing methods, we used five benchmark datasets obtained from Tokheim and Karchin [3] (http://karchinlab.org/data/CHASMplus/Tokheim_Cell_Systems_2019.tar.gz, acquired on October 28, 2021); Kim et al., IARC TP53, Ng et al., Gene panel (OncoKB), and CGC-recurrent.

The hyperparameters of our models used in the evaluation were obtained from the model with the best performance in the five-fold cross-validation. To ensure robust evaluation, we performed this trial three times. Then, three trained models were used to predict each benchmark dataset, and the models were evaluated on the average performance. The prediction performance was evaluated by ROC-AUC and PR-AUC. The thresholds for positive and negative were defined as the average value of the Youden index [24] in the three trained models.

We compared the prediction performance of our approach with 26 preceding tools. There are six tools to predict driver mutations in cancer (CHASM, TransFIC, CanDrA, ParsSNP, CHASMplus, and Network&AA) and there are 20 tools to predict the effect of gene mutations on proteins, not specific to gene mutations in cancer (SIFT [25], MutPred [26], LRT [27], Polyphen2_HVAR [28], Polyphen2_HDIV [28], MutationAssessor [29], PROVEAN [30], VEST4 [31], FATHMM [32], CADD [33], MutationTaster [34], MetaSVM [35], DANN [36], REVEL [37], M-CAP [38], DEOGEN2 [39], MPC [40], ClinPred [41], LIST-S2 [42], and MVP [43]).

We obtained prediction scores of CHASM, TransFIC, CanDrA, and ParsSNP from Tokheim *et al.* [3], Network&AA from Ozturk *et al.*[5], and other prediction scores from the dbNSFP database [44, 45].

### Interpretation of results

We employed SHAP to interpret the contribution of each feature to the prediction results. SHAP is an approach to explain the prediction result, and the SHAP value expresses the contribution of each feature. A larger absolute SHAP value means a larger contribution of the feature to the prediction result. In this study, we evaluated the contribution of the graph features in predicting gene mutations in benchmark datasets using the SHAP values. SHAP Python package (version 0.39.0) was used to calculate the SHAP values.

## Availability of data and materials

The datasets and source codes of Net-DMPred are publicly available at the Github repository, https://github.com/clinfo/Net-DMPred.

Project name: Net-DMPred

Project home page: https://github.com/clinfo/Net-DMPred

Operating system(s): Linux

Programming language: Python

Any restrictions to use by non-academics: No restrictions.

The datasets described in this article can be freely and openly accessed at Pathway Commons (https://www.pathwaycommons.org/archives/PC2/v12/PathwayCommons12.All.hgnc.sif.gz), PathwayMapper (https://www.pathwaymapper.org/) CHASMplus training datasets (http://karchinlab.org/data/CHASMplus/formatted_training_list.txt.gz), five benchmark datasets (http://karchinlab.org/data/CHASMplus/Tokheim_Cell_Systems_2019.tar.gz), and SNVBox (http://karchinlab.org/data/CHASMplus/SNVBox_chasmplus.sql.gz). When running HotMAPS 1D (https://github.com/KarchinLab/probabilistic2020/tree/master), the input datasets were acquired from The Cancer Genome Atlas (TCGA) (https://portal.gdc.cancer.gov/), gene BED file (https://genome.ucsc.edu/cgi-bin/hgTablesa), and gene FASTA file (https://hgdownload.soe.ucsc.edu/goldenPath/hg38/bigZips/hg38.fa.gz).

## Acknowledgments

The analysis in this work were partially performed on the NIG supercomputer at ROIS National Institute of Genetics.

## Author information

### Authors and Affiliations

Graduate School of Medicine and Faculty of Medicine, Kyoto University, Kyoto, Japan.

Narumi Hatano, Mayumi Kamada, Ryosuke Kojima, Yasushi Okuno

RIKEN Center for Computational Science(R-CCS)HPC/HPC- and AI-driven Drug Development Platform Division, Kobe, Japan.

Yasushi Okuno

### Contributions

M.K., R.K. and Y.O. conceived the ideas. R.K.and N.H implemented NetDM-Pred. M.K. and N.H. performed the prediction and analyzed the experiment results. All authors contributed to the preparation of the manuscript.

### Corresponding authors

Correspondence to Mayumi Kamada and Yasushi Okuno

## Ethics declarations

Ethics approval and consent to participate: Not applicable Consent for publication: Not applicableConflict of Interest: None declared.

## Supplementary information

Additional file 1

Supplementary Table 1: ROC-AUC for five benchmark datasets

Additional file 2

Supplementary Table 2:

PR-AUC for five benchmark datasets

Additional file 3

Supplementary Method: The 88 features from SNVBox

