## Supplemental Tables and Method for "Network-based prediction approach for cancer-specific driver missense mutations using a graph neural network"

Supplementary Materials

**Supplementary Table 1.** ROC-AUC for five benchmark datasets.

| | Kim et al. | IARC TP53 | Ng et al. | Gene panel (OncoKB) | CGC-recurrent |
|---|---|---|---|---|---|
| Cancer pathway + Molecular interaction | 0.748 | 0.772 | 0.712 | 0.807 | 0.755 |
| CHASMplus | 0.721 | 0.788 | 0.725 | 0.783 | NaN |
| Molecular interaction | 0.752 | 0.762 | 0.684 | 0.789 | 0.775 |
| Cancer pathway | 0.618 | 0.766 | 0.693 | 0.808 | 0.765 |
| MVP | 0.669 | 0.758 | 0.645 | 0.797 | 0.773 |
| No graph | 0.674 | 0.771 | 0.72 | 0.813 | 0.658 |
| ParsSNP | 0.537 | 0.749 | 0.74 | 0.764 | 0.792 |
| REVEL | 0.581 | 0.78 | 0.645 | 0.819 | 0.748 |
| DEOGEN2 | 0.603 | 0.79 | 0.619 | 0.79 | 0.736 |
| CanDrA | 0.706 | 0.756 | 0.645 | 0.768 | 0.657 |
| M-CAP | 0.676 | 0.739 | 0.554 | 0.762 | 0.785 |
| VEST4 | 0.529 | 0.766 | 0.66 | 0.785 | 0.712 |
| MutPred | 0.533 | 0.773 | 0.656 | 0.799 | 0.672 |
| ClinPred | 0.551 | 0.775 | 0.673 | 0.761 | 0.651 |
| Network&AA | 0.555 | NaN | 0.737 | 0.751 | NaN |
| PROVEAN | 0.551 | 0.77 | 0.654 | 0.756 | 0.577 |
| TransFIC | 0.613 | 0.74 | 0.592 | 0.729 | 0.626 |
| LRT | 0.596 | 0.745 | 0.609 | 0.687 | 0.635 |
| CHASM | 0.691 | 0.688 | 0.578 | 0.645 | NaN |
| Polyphen2_HVAR | 0.46 | 0.76 | 0.614 | 0.732 | 0.672 |
| FATHMM | 0.566 | 0.76 | 0.479 | 0.696 | 0.729 |
| CADD | 0.493 | 0.744 | 0.612 | 0.741 | 0.639 |
| LIST-S2 | 0.559 | 0.714 | 0.636 | 0.673 | 0.635 |
| SIFT | 0.493 | 0.727 | 0.601 | 0.708 | 0.615 |
| MetaSVM | 0.537 | 0.542 | 0.549 | 0.75 | 0.762 |
| MPC | 0.571 | 0.598 | 0.6 | 0.707 | 0.663 |
| MutationTaster | 0.618 | 0.647 | 0.589 | 0.708 | 0.57 |

| | | | | | |
|---|---|---|---|---|---|
| DANN | 0.662 | 0.68 | 0.576 | 0.642 | 0.57 |
| Polyphen2_HDIV | 0.426 | 0.74 | 0.594 | 0.715 | 0.651 |
| MutationAssessor | 0.379 | 0.748 | 0.505 | 0.657 | 0.601 |

**Supplementary Table 2.** PR-AUC for five benchmark datasets.

| | Kim et al. | IARC TP53 | Ng et al. | Gene panel (OncoKB) | CGC-recurrent |
|---|---|---|---|---|---|
| Cancer pathway + Molecular interaction | 0.595 | 0.487 | 0.487 | 0.061 | 0.236 |
| CHASMplus | 0.524 | 0.504 | 0.561 | 0.053 | NaN |
| Molecular interaction | 0.606 | 0.476 | 0.447 | 0.055 | 0.299 |
| Cancer pathway | 0.518 | 0.465 | 0.496 | 0.045 | 0.061 |
| MVP | 0.483 | 0.42 | 0.361 | 0.033 | 0.028 |
| No graph | 0.539 | 0.477 | 0.498 | 0.066 | 0.007 |
| ParsSNP | 0.45 | 0.445 | 0.526 | 0.05 | 0.125 |
| REVEL | 0.347 | 0.461 | 0.343 | 0.036 | 0.058 |
| DEOGEN2 | 0.34 | 0.477 | 0.313 | 0.029 | 0.067 |
| CanDrA | 0.444 | 0.435 | 0.354 | 0.028 | 0.067 |
| M-CAP | 0.554 | 0.378 | 0.278 | 0.023 | 0.04 |
| VEST4 | 0.301 | 0.436 | 0.316 | 0.033 | 0.043 |
| MutPred | 0.421 | 0.426 | 0.391 | 0.037 | 0.021 |
| ClinPred | 0.308 | 0.465 | 0.351 | 0.021 | 0.024 |
| Network&AA | 0.331 | NaN | 0.53 | 0.048 | NaN |
| PROVEAN | 0.307 | 0.426 | 0.328 | 0.019 | 0.042 |
| TransFIC | 0.381 | 0.564 | 0.294 | 0.018 | 0.008 |
| LRT | 0.644 | 0.526 | 0.585 | 0.364 | 0.346 |
| CHASM | 0.651 | 0.35 | 0.351 | 0.019 | NaN |
| Polyphen2_HVAR | 0.343 | 0.49 | 0.374 | 0.161 | 0.153 |
| FATHMM | 0.369 | 0.425 | 0.208 | 0.019 | 0.043 |
| CADD | 0.283 | 0.387 | 0.304 | 0.018 | 0.008 |
| LIST-S2 | 0.357 | 0.369 | 0.326 | 0.017 | 0.008 |
| SIFT | 0.338 | 0.509 | 0.374 | 0.253 | 0.232 |
| MetaSVM | 0.406 | 0.195 | 0.269 | 0.022 | 0.02 |
| MPC | 0.302 | 0.24 | 0.309 | 0.018 | 0.02 |
| MutationTaster | 0.69 | 0.431 | 0.587 | 0.434 | 0.354 |
| DANN | 0.4 | 0.292 | 0.28 | 0.012 | 0.008 |
| Polyphen2_HDIV | 0.317 | 0.525 | 0.453 | 0.304 | 0.274 |
| MutationAssessor | 0.245 | 0.414 | 0.268 | 0.016 | 0.012 |

**Supplementary Method.** The 88 features from SNVBox are below.

AABLOSUM

AACharge

AACOSMIC

AACOSMICvsHapMap

AACOSMICvsSWISSPROT

AAEx

AAGrantham

AAHapMap

AAHGMD2003

AAHydrophobicity

AAMJ

AAPAM250

AAPolarity

AATransition

AATripletFirstDiffProb

AATripletFirstProbMut

AATripletFirstProbWild

AATripletSecondDiffProb

AATripletSecondProbMut

AATripletSecondProbWild

AATripletThirdDiffProb

AATripletThirdProbMut

AATripletThirdProbWild

AAVB

AAVolume

ExonConservation

ExonHapMapSnpDensity

ExonSnpDensity

HMMEntropy

HMMPHC

HMMRelEntropy

MGAEntropy

MGAPHC

MGARelEntropy

PredBFactorF

PredBFactorM
PredBFactorS
PredRSAB
PredRSAE
PredRSAI
PredSSC
PredSSE
PredSSH
PredStabilityH
PredStabilityL
PredStabilityM
RegCompC
RegCompDE
RegCompEntropy
RegCompG
RegCompH
RegCompILVM
RegCompKR
RegCompNormEntropy
RegCompP
RegCompQ
RegCompWYF
UniprotACTSITE
UniprotBINDING
UniprotCABIND
UniprotCARBOHYD
UniprotCOMPBIAS
UniprotDISULFID
UniprotDNABIND
UniprotDOM_Chrom
UniprotDOM_LOC
UniprotDOM_MMBRBD
UniprotDOM_PostModEnz
UniprotDOM_PostModRec
UniprotDOM_PPI
UniprotDOM_RNABD

UniprotDOM_TF
UniprotLIPID
UniprotMETAL
UniprotMODRES
UniprotMOTIF
UniprotNPBIND
UniprotPROPEP
UniprotREGIONS
UniprotREP
UniprotSECYS
UniprotSIGNAL
UniprotSITE
UniprotTRANSMEM
UniprotZNFINGER
InsiderPPI
NumAlignedSpecies
UniprotDensity
